# Schooling of Light Reflecting Fish

**DOI:** 10.1101/2022.12.13.520208

**Authors:** Assaf Pertzelan, Gil Ariel, Moshe Kiflawi

## Abstract

One of the hallmarks of the collective movement of large schools of pelagic fish are waves of shimmering flashes that propagate across the school, usually following an attack by a predator. Such flashes arise when sunlight is reflected off the specular (mirror-like) skin that characterizes many pelagic fishes, where it is otherwise thought to offer a means for camouflage in open waters. While it has been suggested that these ‘shimmering waves’ are a visual manifestation of the synchronized escape response of the fish, the phenomenon has been regarded only as an artifact of esthetic curiosity. In this study we apply agent-based simulations and deep learning techniques to show that, in fact, shimmering waves contain information on the behavioral dynamics of the school. Our analyses are based on a model that combines basic rules of collective motion and the propagation of light beams in the ocean, as they hit and reflect off the moving fish. We use the resulting reflection patterns to infer the essential dynamics and inter-individual interactions which are necessary to generate shimmering waves. Using an artificial neural network, trained to classify the direction of attack and the shape of the school based only on the flashes, we also provide a proof-of-concept, showing that flash patterns are indeed indicative of the state and dynamics of the school and the individuals composing it. Moreover, we show that light flashes observed by the school members themselves extends the range at which information can be communicated across the school. To the extent that the fish pay heed to this information, for example by entering an apprehensive state of reduced response-time, our analysis suggests that it may speed up the propagation of information across the school.

## 1. Introduction

Free-ranging animals that move in large aggregates, such as schooling fish and flocking birds, are often required to maneuver in unison to evade attacking predators. The collective evasive maneuvers change macroscopic properties of the aggregate, such as its direction, polarization, and density (Radakov 1973; Gerlotto et al. 2006; Hemelrijk et al. 2015; Herbert-Read et al. 2015). These changes are often manifested as “waves of agitation” or “escape waves” that follow the propagation of information regarding the attack across the aggregate. The complex inter-individual interactions, which bring about such large-scale dynamical patterns, are not fully understood (Gerlotto et al. 2006; Hemelrijk et al. 2015; Herbert-Read et al. 2015). One of the main empirical obstacles to a fuller understanding of these interactions and the consequent escape dynamics is the difficulty of tracking individuals within very large schools; using either cameras (e.g. (Brehmer et al. 2019)) or sonars (Gerlotto et al. 2006; Brehmer et al. 2019).

Large schools of silvery fish are common in the world’s oceans (Denton and Nicol 1966). When swimming close to the surface, the specular (mirror-like) skins of these fish will often reflect direct sunlight. To an underwater observer, these reflections appear as highly conspicuous flashes of light (Figure 1), which increase the fish’s contrast by at least one order of magnitude (Pertzelan et al. 2022). When under attack, an agitation wave crossing a school of specular fish, may appear as a shimmering wave of flashes (e.g. Figure 1). The wave arises as the succession of evasive maneuvers by neighboring school members momentarily brings their bodies to an angle that reflects the sun in the direction of the observer (Radakov 1973; Gerlotto et al. 2006; Hemelrijk et al. 2015; Herbert-Read et al. 2015). As such, shimmering waves could contain information regarding the dynamics of the school and individuals within it, which is more discernable than the actual trajectories of the individual fish.

**Figure 1.**
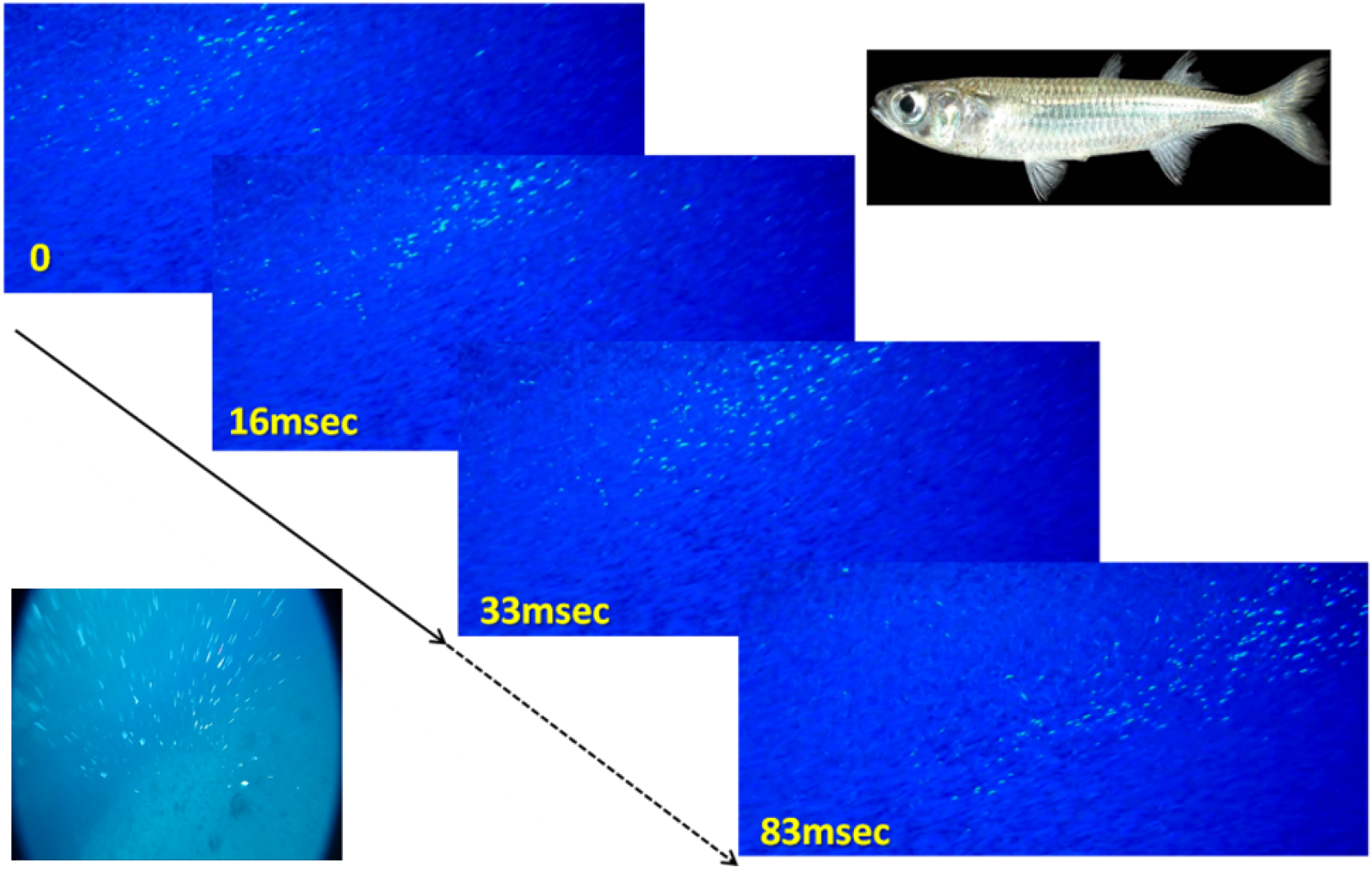
Propagation of an agitation (shimmering) wave in a school of *A. lacunosus* (top-right corner), filmed at 60 frames per second. The flashes arise when the fish, initially facing to the left, turn into the school (here into the page, perpendicular to the advancing wave), causing their body to face the observer at just the right angle to generate a flash of reflected sunlight. The wave propagates as the decision to turn is taken sequentially by the fish, starting from the left. Bottom-left corner: a “predator’s view” of the wave, taken from a camera mounted on a spear shot into the school.

In this paper, we use individual-based simulations to study the dynamics of shimmering-waves. We begin by testing whether a shimmering wave can arise in possible schooling scenarios. We then test the complementary question of whether the dynamic properties of the waves (e.g., their duration and speed of propagation) can be used to differentiate between competing hypotheses relating to the inter-individual interactions that give rise to them; as well as to infer the occurrence and direction of an attack. Finally, we test the possibility that flashes observed by the fish themselves can be used to induce apprehension and, thereby, speed-up the transfer of information within the school.

### Fish schools under attack

In schooling fish, the response to an attack amounts to moving closer together or rolling sideways (Radakov 1973; Gerlotto et al. 2006; Hemelrijk et al. 2015). Once initiated, waves of agitation often propagate faster than the speed at which individuals are moving within the group; a phenomenon often referred to as the ‘Trafalgar effect’ (Treherne and Foster 1981). In cases where the stimulus is an approaching predator, the waves have also been found to travel faster than the predator itself (Herbert-Read et al. 2015). Surprisingly, waves can travel even faster than expected given the estimated response-latencies (Hemelrijk et al. 2015), i.e., the time between the perception of the stimulus and the fear/escape response. The unexpected speed of information-transfer led to speculations regarding possible mechanisms that enhance synchrony and extend beyond the scale of localized social interactions. A prominent example is the “chorus-line hypothesis” (Potts 1984), which assumes a reduction in latency due to a heightened state of anticipation.

Importantly, the flashes of light reflected off specular fish are visible not only from outside the school but also from within it; and thus, could potentially serve to inform the school members themselves. Indeed, it has been previously hypothesized that the reflective structures on fish can be used for “communicating information on relative positions, orientations, and movements between neighbors” (Denton and Rowe 1994). Here, we explore a hypothesis that the heightened state of anticipation described above is caused by changes in the pattern of reflected light, as perceived by school members found downstream of the propagating wave. Particularly, we propose that observing a large change in the number of flashes reduce the latency and, as a result, speed-up the propagation of information.

### Modelling schools under attack

Standard models of schooling typically fall short of generating agitation waves following localized perturbations, such as an attacking predator; presumably because the perturbation is perceived by only a small number of agents (Herbert-Read et al. 2015; Sonoda et al. 2019). In most models (for example the three-zones model e.g. (Aoki 1982; Reynolds 1987; Vicsek and Zafeiris 2012)), individuals modify their position and direction based on the average response of neighboring agents. As a result, the initial reaction to the perturbation is “averaged out” (Radakov 1973; Gerlotto et al. 2006; Hemelrijk et al. 2015; Sonoda et al. 2019), i.e., the response of an agent close to the perturbation is averaged with agents that are farther away and, thus, did not perceive it. As most agents are far from the perturbation, the intensity of the reaction decreases with distance. Indeed, subsequent models have shown that additional responses, that are not due to a direct observation of the predator but leads to preemptive evasive maneuvers, are needed in order to generate agitation waves (Hemelrijk et al. 2015; Sonoda et al. 2019). For example, (Sonoda et al. 2019) assumed that when an agent sees another agent whose behavior is clearly different from other school members, it will copy its behavior. With this added behavioral component, information of an approaching predator can propagate through the school, forming a response wave that travels at a (approximately) constant speed (Herbert-Read et al. 2015). The agitation wave gives rise to a collective evasive response, even though the number of individuals that directly experience the perturbation is small (Sonoda et al. 2019).

The simulations introduced below accommodate the three basic local-interaction rules which are typically included in collective-motion models (repulsion, attraction, and alignment), as well as the copy response suggested in (Sonoda et al. 2019). On this collective-motion model we superimpose a ray-tracing model (Pertzelan et al. 2022) that ‘records’ the light-flashes perceived from a pre-prescribed location, either within or outside the school. To the best of our knowledge, our model is the first attempt to model shimmering waves, based on first principles of light propagation and reflectance.

## 1. Methods

Our modeling strategy is a second-order agent-based model (i.e., we specify the acceleration). Simulations include a school of fish and a single predator. The latter appears for a limited number of steps and moves along a fixed linear trajectory. The predator moves towards and into the school faster than the agents’ maximal speed during the ’schooling’ state (see below).

In the three-zones model (e.g (Aoki 1982; Reynolds 1987)), each agent *i* at every discrete time-step *t* is described by its position, *p_i_*(*t*), and velocity, *v_i_*(*t*). To account for predator response (the *direct-only* model), we introduce two additional variables: the acceleration *a_i_*(*t*), and a discrete internal state *s_i_*(*t*). The internal state is one of four options: schooling, evasive, predator-response (similar to (Lee 2006)), or copy-response (similar to (Sonoda et al. 2019)), as described below. Moreover, in order to trace the direction at which light-rays reflect off the fish, we keep track of the vertical direction of each agent. To this end, we define a direction *b_i_*(*t*) that is perpendicular to the velocity, corresponding to the back of the fish (the direction that regularly points upward with the vertical axis).

Below, we outline the main components of the model. Details are provided in the SI section 1.

### Agent dynamics

At each step, all agents update their positions and velocities synchronously as,

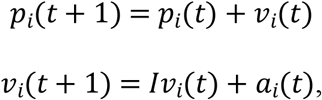

where *I* < 1 is an inertia parameter, qualitatively accounting for water resistance. Velocity-updates depend on the current internal state *s_i_*(*t*), according to the decision process explained below and depicted in Figure 2.

**Figure 2.**
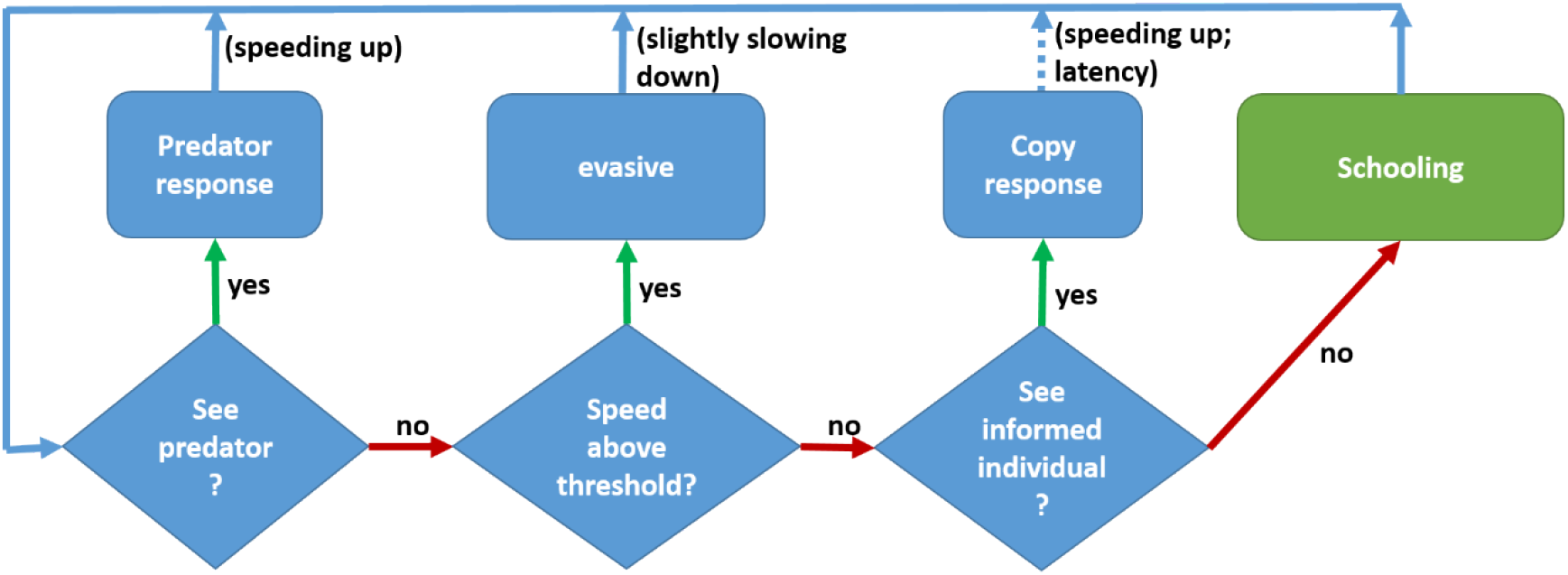
State diagram. A diagram of the decision process for moving between states, *s_i_*(*t*). The states are represented by rectangles and the conditions by diamonds. If an agent sees a predator it accelerate to a high speed turning away of the predator. If an agent sees an informed individual (a high-speed individual facing away of the agent direction), it copies this informed individual velocity. From then on, it remains in the *evasive* state until it gradually slows down and return to the *schooling* state. The copy response occurs after several steps of latency (default: 2 steps; corresponding to 0.084 msec. see Table 1), during which the agent remains in the *schooling* state.

See Table 1 for conversion of the simulation parameters to empirical estimates.

**Table 1.**
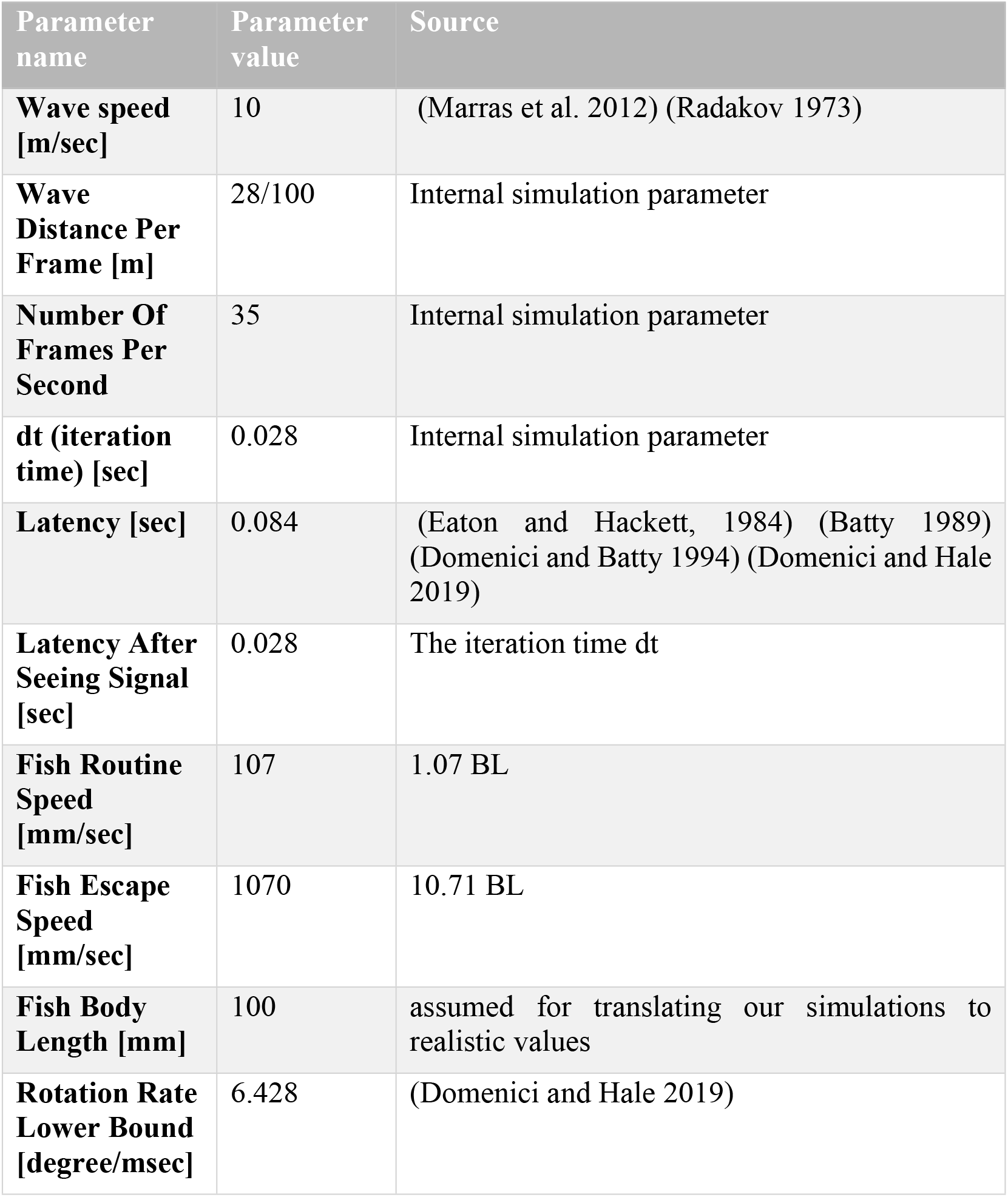
Translation of the simulation parameters to real-world values.

### Internal states

- *Schooling*: An agent’s acceleration is determined similar to the usual three-zone rules, e.g. (Couzin et al. 2002), i.e. a weighted average of the attraction to all neighbors within a given distance *R*_o_, alignment with all neighbors within a distance *R*_a_ and repulsion from neighbors within a distance *R*_r_; with *R*_r_<*R*_a_<*R*_o_ (Figure S1).
- *Evasive*: If an agent in the *schooling* state attains a speed that is higher than the schooling speed limit 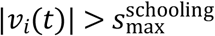 (for example, because it encountered a predator or an informed individual – see *copy-response* below), it transitions to an *evasive* state. In this state, the update rules for the velocity are similar to rules while *schooling*, but with different weights (Table S1) that give a higher priority to cohesion and alignment over repulsion.
- *predator-response*: If an agent in the *schooling* or *evasive* states encounters the predator (i.e., the predator is within a distance *R*_p_), it transitions into the *predator-response* state. In this state, agents turn towards the opposite direction of the predator and start swimming at speed 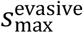.
- *copy-response*: A neighbor (to any *schooling* agent within radius *R*_i_) is called *informed* if it is currently making a sharp turn and is swimming fast (above a threshold). Such a behavior indicates (possible) information of an attack. A *schooling* agent that has an informed neighbor will transition into the *copy-response* state following a response delay (latency). In this state, agents copy the velocity of the informed neighbor. In the following, we set *R*_i_ = *R*_a_, as both represent the distance at which the orientation of neighbors are visible.

#### Roll angles

In order to track the angle at which incident light beams are reflected off the agents, we need to model the dynamics of a direction normal to the velocity, corresponding to the ventral side of the fish. We denote this direction as *b_i_*(*t*). To this end, we define a motion rule for the roll-alignment, which determines the torsion of the trajectory curve of each agent. In addition, we give individuals a tendency to realign *b_i_*(*t*) with the vertical axis (point upwards). See sections 1 in the SI for details.

#### Optics

We assume a light source above the water-surface that is a cone of light of angle *θ* around an incident direction *d*_light_. In all simulations, we take *d*_light_ = – *i*_3_ = (0,0, – 1), i.e., pointing directly downwards. For simplicity, the agents are considered as 2-sided planar mirrors whose normal direction *n_i_*(*t*) is perpendicular to the velocity and back vectors, 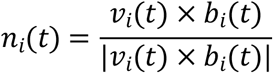. A more detailed model that provides a realistic representation of light reflected off the skins of silvery fish is given in (Pertzelan et al. 2022). This model, which is computationally expensive, was only used for static snapshots of fish schools with no dynamics. In order to determine if agent *i*. reflects light towards position *x* in a given time *t*, we define a vector in the direction of *p_i_*(*t*) – *d*_light_, 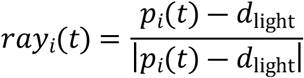. Then, we define the reflected vector of *ray_i_*(*t*) from the plane represented by the normal *n_i_*(*t*),

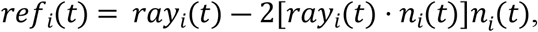

and another vector pointing from the agent position towards a virtual observer that is placed at position *x*,

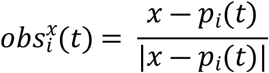

If the angle between *ref_i_*(*t*) and the vector 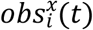 is smaller than *θ*, then the agent reflects light towards *x*. It is possible that an agent will not reflect light both at time *t* and at time *t* +1, but its trajectory for its location between the two times will cross a point in which it will flash. This is taken into account, as explained in the SI (section 1, figure S3).

#### Classification of behavioral patterns

To differentiate between some of the different behavioral and structural configurations accommodated by the model, we apply an artificial deep neural network (DNN) to the ‘recorded’ pattern of reflections. DNNs are increasingly used in the study of collective animal behavior (Hughey et al. 2018; Dunn et al. 2019; Jeckel et al. 2019; Raghunathan 2021; Achuthan et al. 2022), owing to their capacity to find features that are intuitive to recognize but unintuitive to define. In particular, DNNs are efficient in detecting features in images taken from different perspectives and scales (Long et al. 2015) and in sequences of inputs (Shi et al. 2015; Yin et al. 2020; Jagtap and Chavaan 2021). Since the training of DNNs requires large, annotated datasets, researchers are increasingly turning to synthetic data (Mitash et al. 2017; Xu and Goodacre 2018; Dunn et al. 2019; Raghunathan 2021; Achuthan et al. 2022). We adopt a similar approach, using our model to facilitated supervised leaning by the neural network.

Below we present three sections, each combining the technical details and results for each one of our main objectives. First (section 3), we demonstrate that the model successfully produces shimmering waves using realistic parameter values (Table 1); contingent on the inclusion of a copy response, i.e. consistent with (Sonoda et al. 2019). Secondly, (section 4) we demonstrate the ability to distinguish between different characteristics of the school, based only on observed flash patterns. Third (section 5), we test the hypothesis that the unique flash signature caused by an attack can: 1) be seen by fish found downstream from the propagating wave and, were the fish to use this information, 2) leave a discernable signature in the dynamics of the waves.

## 2. Generation of flash waves

Our modeling approach enables us to demonstrate that a model implementing both *predator-response* and *copy-response* can successfully generate seemingly realistic shimmering waves. Moreover, the copy response is necessary, i.e., the basic three-zone interaction rules and *predator-response* alone are not sufficient to explain the waves seen in nature.

### Methods

All schools are composed of *N*=15,000 agents, initially positioned in an elongated cylinder parallel to the x-axis (assuming a body length of 10cm and a density of ~33 fish/m^3^, an average distance of ~40cm between agents, corresponding to a cylinder of around 28m long and 5.5m wide; see SI table 1 for the full configuration).

A virtual observer was placed~12.5m from the school center of mass on the same horizontal plane and perpendicular to the school average velocity, i.e., the observer is watching the school from its side (Figure 3). During each scenario, the observer remains at the same relative position to the center of the school (that is practically stationary) but not moving relative to the orientation of the school. The predator attacks the school head-on and aims its attack at the center of mass of the first 30 individuals, evaluated in relation to the average velocity of the school. The predator was removed from the system at the end of the attack, which lasts five time-steps. Simulations resolve the group dynamics for 115 simulation steps, corresponding to approximately 4 seconds (see Table 1). See SI sections 12-15 for sensitivity of the appearance of the wave.

**Figure 3.**
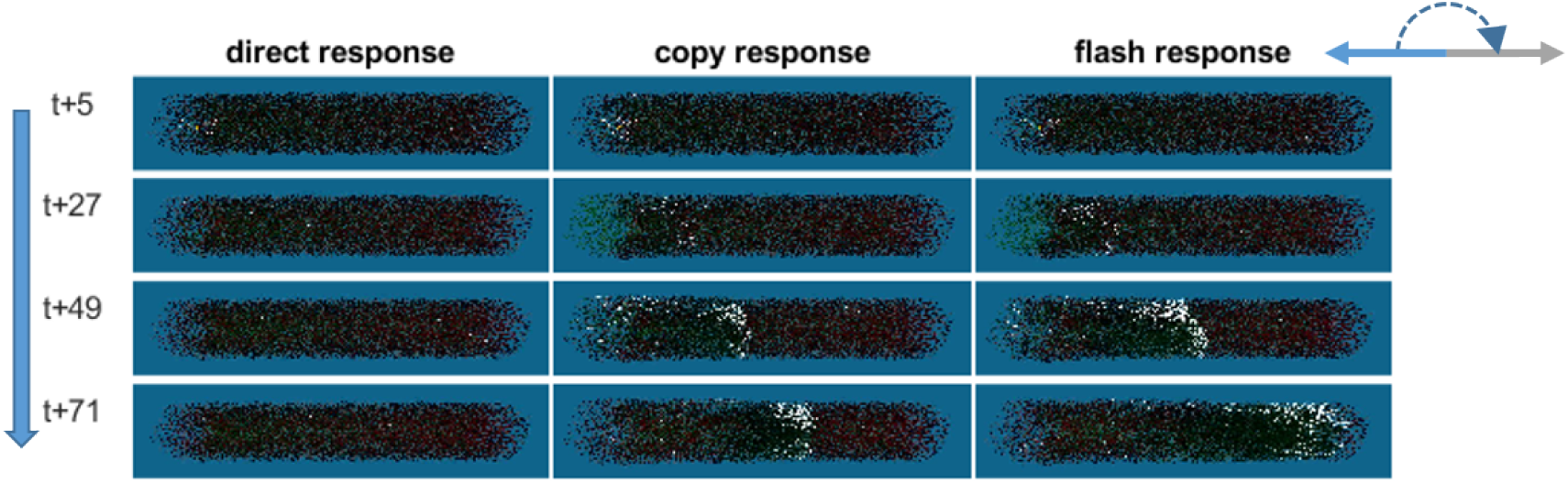
Snapshots of the dynamics that develop following a head-on attack on a cylindrical school swimming to the left, under each of the three models (columns). Time-steps are shown on the left, and progress from top to bottom. The shimmering wave appears as fish reflect light (turn ‘white’) toward the observer, as they turn approximately 180 degrees (arrows at top-right corner) to face the other direction and evade the predator.

We compared two response models (Figure 3, left and middle columns) that differ in their velocity-update mechanisms:

- *Direct-only*: A fish in the schooling state that encounters the predator directly performs an evasive response (that increases their speed, see Section 2). Under this model, the fish react only to a direct observation of the predator.
- *Direct-and-copy*: In addition to the direct response, fish found in the schooling state which did not see the predator, but do see an informed individual, enter a *copy-response* state and copies the velocity of the informed agent. Copying occurs with a latency of 3 steps (Table 1), corresponding to the reaction time of a fish.

To quantify the dynamics that follow the attack, under each of the two models, we first divided the school into ~300 bins (slices) along the x-axis, each corresponding distance of 100mm. For each bin, at each time-step, we then calculated three measures:

a. Change in local fish density: The number of fish in bin *i* at time *t*, expressed in standard deviations (z-score, using the mean and variance calculated across time-steps). As in previous works (Hemelrijk et al. 2015; Herbert-Read et al. 2015; Lecheval et al. 2018), we expected a wave of temporarily increased numbers (i.e., density) to propagate across the school.
b. The proportion of fish that were perceived as flashing by the external observer. This is relevant for establishing the emergence of shimmering waves as a result of escape maneuvers.
c. The proportion of fish that encounter at least one informed individual, at time *t*. This measure quantifies the actual propagation of information.

Each of these measures was plotted, using a heatmap as a function of both the position along the *x*-axis of the school and time. Patterns in these heatmaps were used for inferring the dynamics of the respective measure. In particular, the presence of a diagonal in a heatmap was taken as indicative of the propagation of the respective measure, with the slope of the diagonal equaling the reciprocal of the speed of propagation.

### Results

As expected, a localized predator attack under the *direct-only* response model did not produce a shimmering wave (Figure 3, left column), indicating a rapid loss of information due to averaging-out of the evasive response. Indeed, all measurements under the *direct-only* model do not reveal any meaningful patterns or indicate propagation of information (See SI section 8 for the plots).

In contrast, the same attack under the *direct-and-copy* response model produced the expected shimmering wave, which traversed the school at an approximately constant speed (Figure 3, middle column). Neighbors of informed individuals copy its behavior, without loss of the information regarding the direction of the attack. As a result, the copied response propagates through the entire school within 115 steps.

The *direct-and-copy* model produced a density wave, which appears as a diagonal in Figure 4a. The propagation of the density wave is matched by the pattern of flashes perceived by the external observer (Figure 4b), which follows the actual propagation of information across the school (Figure 4c). Thus, the rate at which fish were exposed to an informed individual (and copied its behavior) matches the density and shimmering waves.

**Figure 4.**
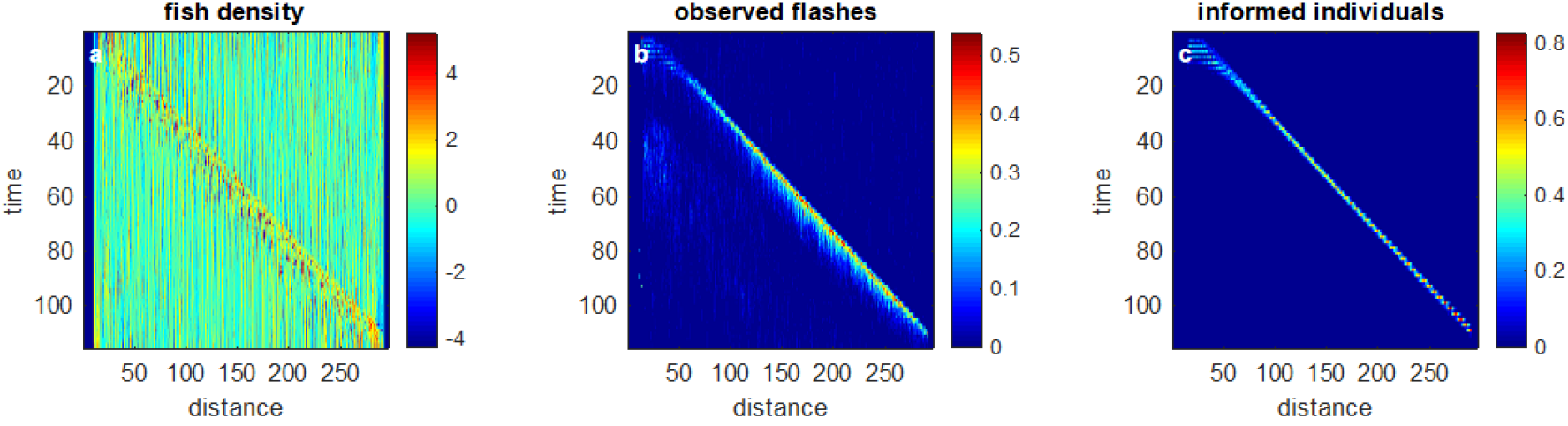
The *direct-and-copy* model, with latency of 3 time-steps. For each plot, the x-axis represents a spatial division of 3D-space along the x-axis, which corresponds to the length of the school. The y-axis represents the timeline, starting with the attack at t=0 and running for a total of 115 steps. a A wave of changes in density(z-score) progresses with a constant speed along the school. The leftmost part (*x, t*=0) is distorted due to direct predator responses. b. The proportion of flashing fish in each bin, as seen by the external observer. The flash pattern that is visible to the observer is consistent with the density wave. c. The actual propagation of information, i.e. the rate of new fish that were exposed to an informed individual and copied its behavior. The slope of the diagonal represents the speed of the wave.

As expected, sensitivity analysis showed that: wave speed decreases with latency (Figure S14a) and increases with escape speed (the initial evasive speed; Figure S14b). The wave-speed dependence on distances of all the motion rules (Figure S14c) and the distance at which the fish applies the *copy-response* (Figure S14d), is approximately linear.

## 3. The information content of shimmering waves

The previous section describes an agent-based model that recreates shimmering waves as a response to predator attacks. The goal was to generate flash patterns that resemble realistic ones. Here, we take a complementary approach and demonstrate that the flash patterns can be used to differentiate between different scenarios relating to the schooling fish. In other words, we aim to show that such differentiation is indeed possible rather than look for the features that make it possible. As proof-of-concept, we focused on two analytical tasks: the detection of an attack and its direction, and the inference of the shape of the school under attack (Figure 5).

**Figure 5.**
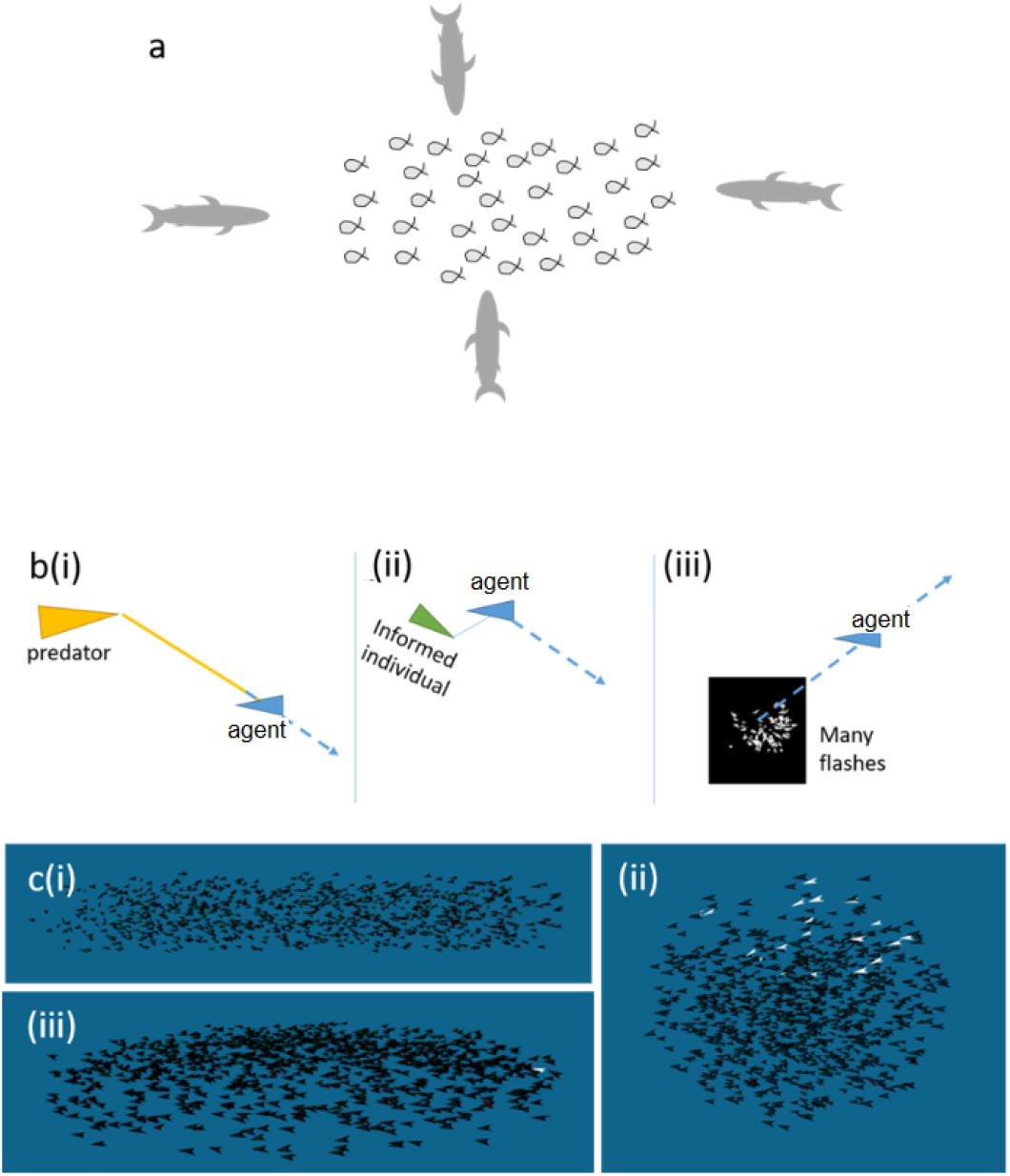
Classification consists of three parameters. a. Attack direction: front, back, top, bottom, side, and no-attack. b. Response-mechanisms: (i) direct-only (ii) direct-and-copy response (iii) flash-direct c. School shape before the attack was initiated: cylindrical, ball, and ‘pancake’ shaped (*N*=1000 fish).

### Methods

#### Task 1. Detect an attack and its direction

We simulated a predator attacking a school of fish and the consequent response of the school. We created six classes corresponding to attacks from five different directions: front, back, side, top, and bottom, and an additional “no-attack” scenario (Figure 5a).

#### Task 2. Infer the shape of the school

We compared the flash signature of attacks on three different school shapes: elongated cylinder, ball, and pancake (a thin circular school close to the water surface). In all simulations, school size was *N*=1000. At the beginning of each simulation, all the fish were oriented in same direction (Figure 5c).

For each class of each task, we generated multiple samples – i.e., short, simulated videos. For each sample we recorded only the flashes, not the ‘fish’ themselves, and saved a tag specifying the corresponding class. See SI section 3 for an example.

##### Datasets generation

For each of the two tasks we generated 150 samples per class using the *direct-and-copy* response model described above. Each simulated sample consisted of a short image-sequence (a ‘video’) of 15 frames, 90×90 pixels each (equivalent to 0.2-0.8 seconds). See, for example, Figure S5, showing only the flashes that are visible to the observer.

In order to differentiate between the classes, we used a convolutional-LSTM network (Based on (Shi et al. 2015). Intuitively speaking, convolutional networks are efficient for detecting variations of image features regardless of scale and position of the camera. LSTMs are useful for processing sequences by adding a time-dimension to the process and tying together features of frames in different times, based either on timespan or on a set of features that were detected in between. In our case, when the observer neither knows its exact position nor his movement in relation to the group, and when there is a strong temporal factor, the combination of convolutional and LSTM architectures is appropriate (Shi et al. 2015; Yin et al. 2020; Jagtap and Chavaan 2021).

##### Our model

The model was written in Python using Keras and was trained and tested on the platform of Google Colabs. Our model consists of a ConvLSTM2D layer which expects tagged sequences of 90×90 grayscale images. A dropout of 0.2 was added to this layer. This layer is made of 64 convLSTM cells each and applies 5×5 convolution filters as their gates and input/output. The output of this layer is transformed to be one-dimensional and is transferred to a fully connected layer (dropout: 0.3). The nodes in this layer are Rectified linear units (ReLU). Fully connected ReLU layers are often used as default layers because of performance considerations during training (Brownlee 2019). This ReLU layer is fully connected to our output layer, which consists of nodes at the same number of classes in our dataset that apply softMax on the input. The softMax function is normalizing the inputs into probabilities for each class with amplification of the probabilities of the higher input values. See SI for the code and its description. Training: the data was divided approximately 4/5 for training (~120 samples per class in our case) by 40 epochs of batch size 8. The remaining 1/5 of the dataset (~30 samples per class in our case) was left for testing. A patience of 7 was added to avoid overfitting.

##### Evaluation of the model

To analyze the quality of a trained model with the test datasets we used confusion matrices that compare the frequency of true and the predicted classes (~30 samples per class). From the confusion matrices we calculated the accuracy, precision, recall and f1-scores.

##### Random noise

To test the robustness of the classification to missing information, we added three (optional) uncertainty parameters (figure S6). These parameters were set to reflect information that is not available to the observer in realistic situations: a. *roll-noise*: The degree of random roll of the fish around its head-tail axis, b. *lookat-moves*: The observer’s position could stay relatively fixed but the direction of the center of its field of view may move (for example, near a wavy sea surface), and c. *observer-orientation*: An observer can tell where it stands in relation to the school on the vertical plane (if it is looking up or down) but not in the horizontal plane.

### Results

Below we demonstrate our ability to infer the state of the school under each of the following scenarios, given that light flashes are the only information available.

The results were derived from analyzing the ‘noisy’ datasets, generated by models that included ‘observer-orientation’, ‘look-at-moves’, and ‘roll-noise’. See the SI section 3 for results without noise.

### Attack-direction

The trained network can detect with high accuracy whether an attack occurred or not (f1-score of 0.92 for the no-attack scenario), and can detect, with a lower accuracy the attack direction. With all noise parameters on, the overall accuracy of this model is 0.74 which is better than chance (1/6), but indicates that there are still some difficulties in distinguishing between the classes (See SI section 5.3). These difficulties could be due to either essential similarities between the flash signatures, or non-optimal selection of hyperparameters (Figure 6).

**Figure 6.**
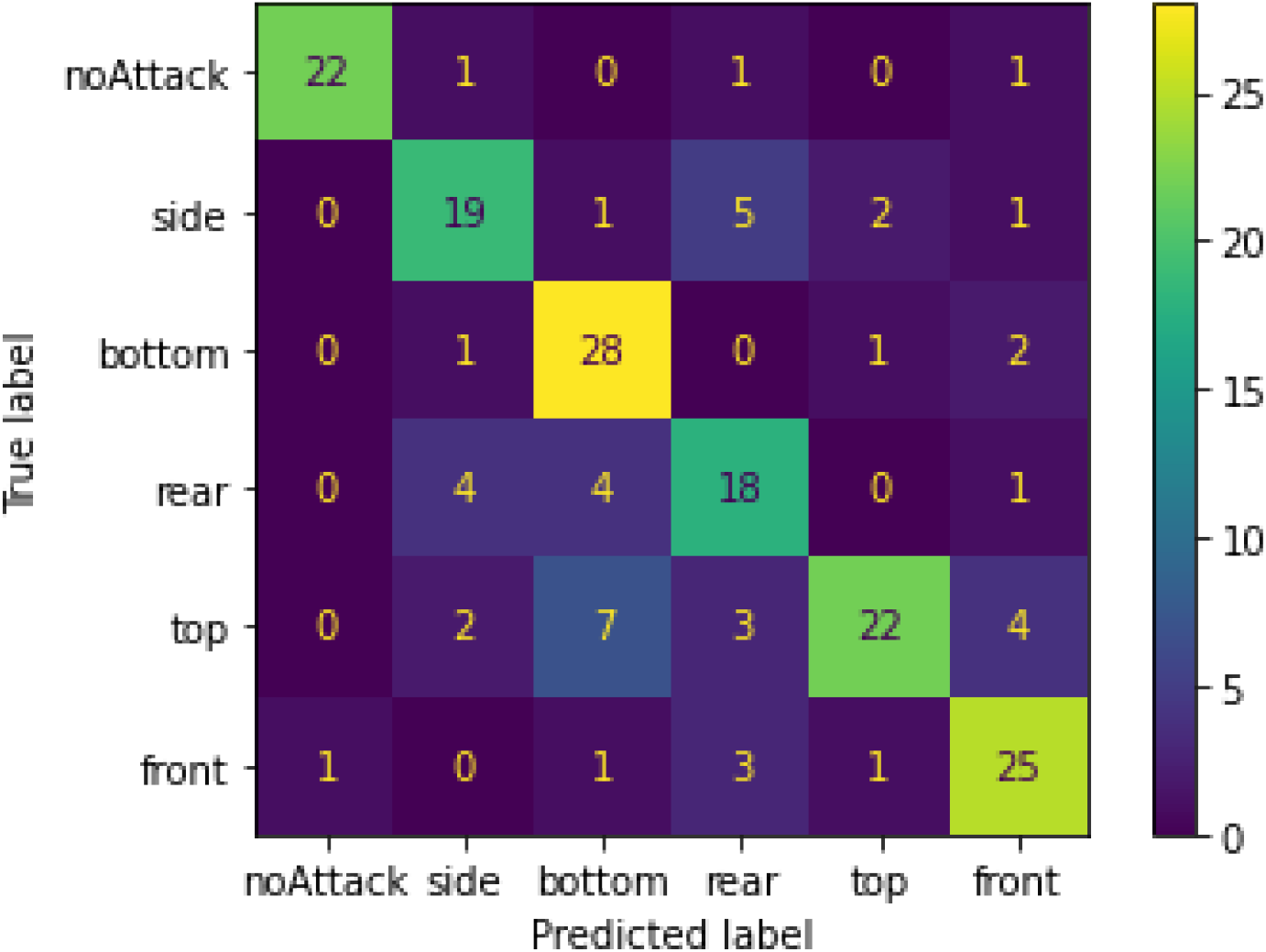
Confusion matrix for ‘attack-direction’. The labels related to the attack directions relatively to the fish heads (represented in the simulation by the velocity vector). The DNN could distinguish among different attach direction in most cases.

#### School shape

The shape of the school had higher accuracy (0.94, See SI section 5.3). This level of accuracy would aid the differentiation between other properties (e.g., attack-direction) when the shape of the school is unknown, as the school-shape could easily be filtered out. It should be noted that this level of accuracy pertains to the school-size and observer-distance specified in the model, and may change if these parameters are changed (Figure 7).

**Figure 7.**
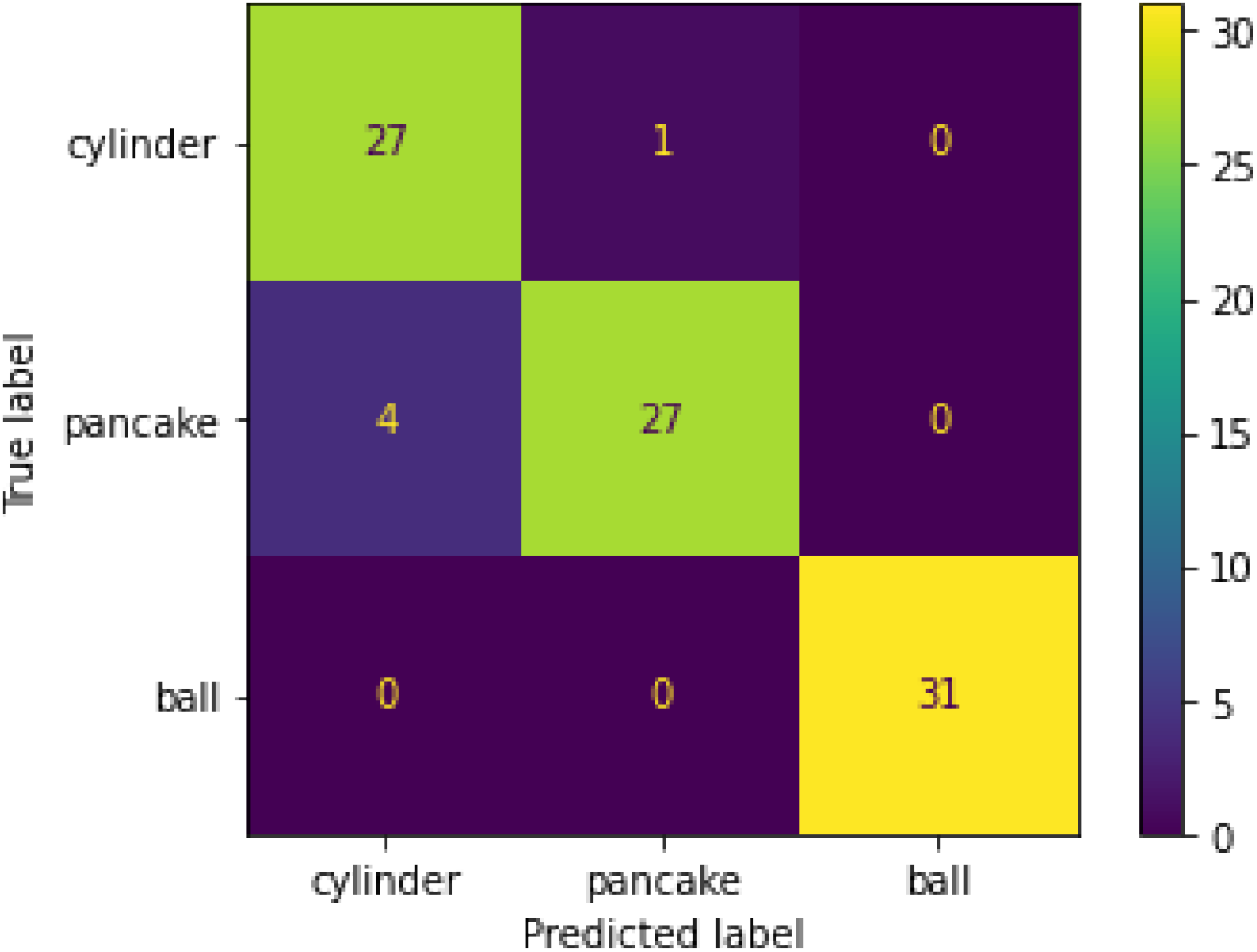
Confusion matrix for ‘school shape’. The DNN could distinguish among different school shapes in most cases. Some pancake-shaped schools have been mistaken for cylinders, possibly due to sharing a large school depths from certain angles.

## 4. Light flashes as a mean to speed up information transfer

In this section we introduce a new assumption – namely, that fish within the school perceive and act upon changes in the number of flashes. To incorporate a ‘flash response’, we assume that observing strong shimmering (the turning on or off of a large number of flashes, per time unit) is a signal for instability in the motion of the school. We further assume that agents respond to this signal and, as a precaution to a possible attack, will enter an anticipation mode that shortens their response-latency and, thereby, results in a faster evasive maneuver.

Given the high detectability of the flashes, the number of flash changes is visible in all areas of the school, including those through which the propagating wave of informed individuals has not yet reached. As a result, the wave can be used as an early warning. Figure 8 demonstrates how the flash signal is viewed by a virtual fish at the back of the school.

**Figure 8.**
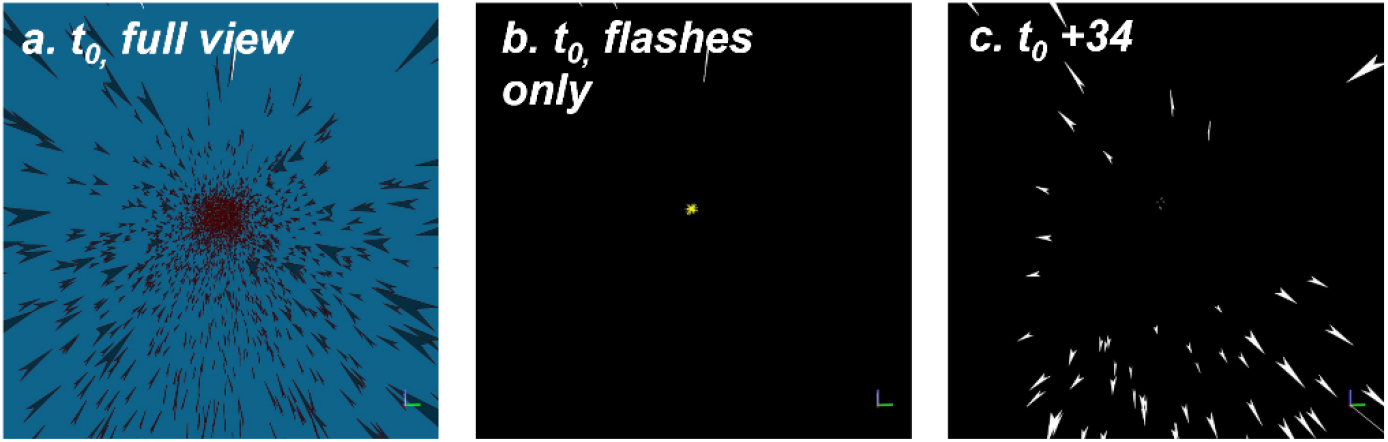
The view from the back of the school (N=5000). ‘*t*_0_, full view’ shows the full-detailed view of the school, one time step before the attack. This view, which shows the fish rather than the flashes, cannot be fully seen by the fish due to the low contrast of the non-flashing school members. Plots b shows the same figure with the few high-contrast flashes that are visible to the observer, and the predator in the distance (in the middle of the frame). After 34 steps (plot c), the flash wave gets closed to the observing fish. For illustrative purposes we chose the fish to be at the “back” of the school and present a narrower field of view than that which is available to the simulated fish. For clarity, and as in the tested models, we ignore the occlusion of flashes. See the section 7 in the SI for the lack of effect of occlusions in such scenario.

### Methods

We modified the *direct-and-copy* response model by specifying how a fish respond to an abrupt change (increase or decrease) in the number of flashes, which falls above some specified threshold (+/- 200): (i)*flash-direct*: the fish turns away from the center of mass of those changes; (ii)*flash-latency*: the fish reduces, to zero, the latency of its response.

Next, we trained the DNN, on the (simulated) flash patterns registered by an external observer, to see if it can distinguish between the original response models (direct-only and direct-and-copy) and the two modifications (flash-direct and flash-latency). In other words, we wanted to establish whether the use of light flashes as a precautionary signal by the fish, will leave a discernable signal in the shimmering wave generated by a collective evasive maneuver.

### Results

With the flash-latency model, the dynamics is initially similar to the one with the *direct-and-copy* response model (Figure 3, right column). However, at some point (~*t*=40) the fish downstream of the propagating wave perceive an above-threshold change in the overall flashing pattern which shortens their latency and consequently, speed up the propagation of the wave (Figure 9**Figure 9**).

**Figure 9.**
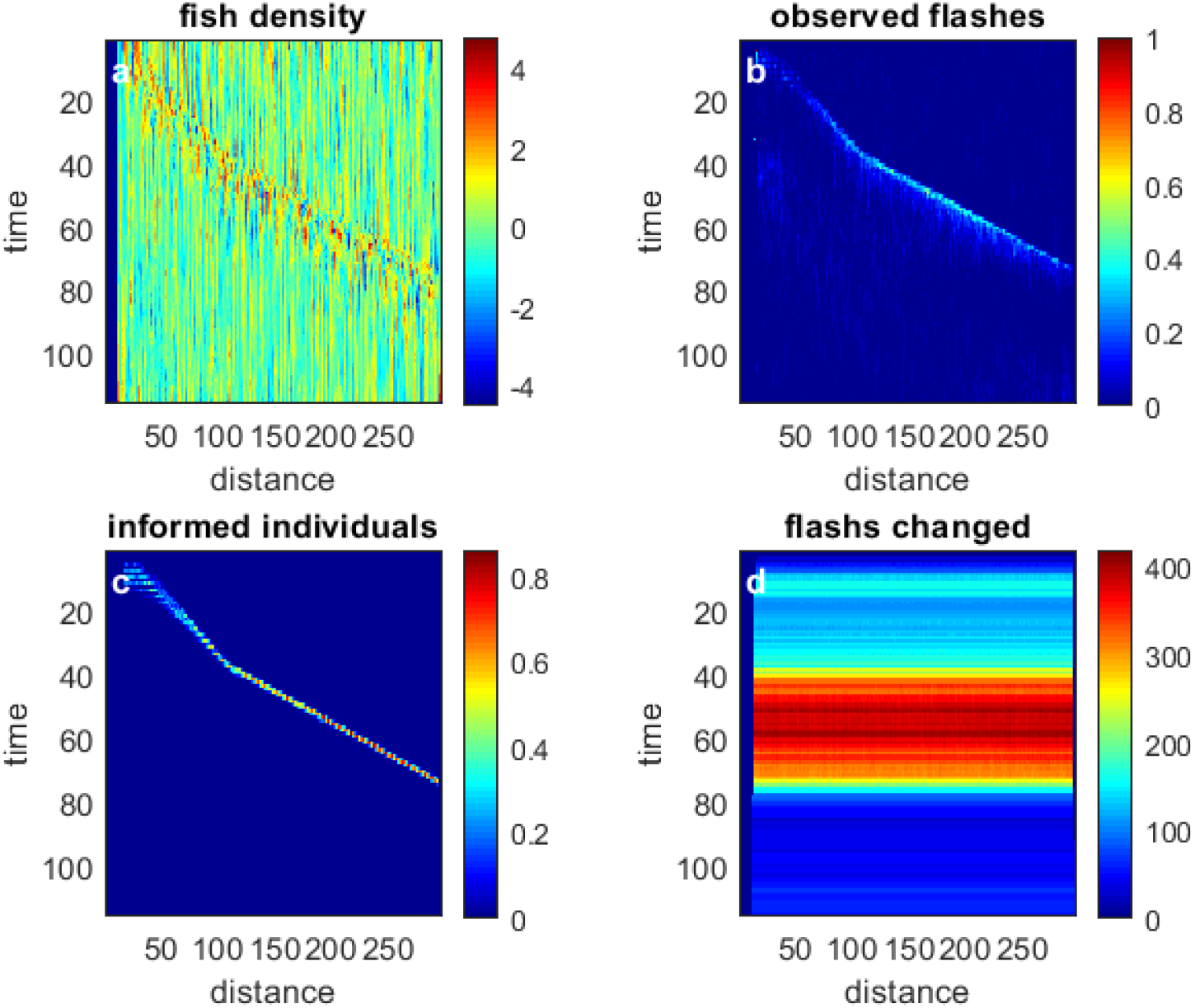
The four basic measures for the *flash-latency* model. The wave starts at the same speed as the copy-response scenario (compare plots a, b, and c to their corresponding plots in Figure 4) but accelerates around *t*=40. At this time, the flash signal crosses a threshold (set to 200; plot 4d) for all fish, which reduces the response latency, accelerating the propagation of information.

The accuracy (see table S2 for the measurements of the DNN) of this classification was 0.74 (expected 0.25 by chance). The identification of *direct-and-copy* and the *flash-latency* scenarios is good, and only slightly mixing with each other (f1-score of 0.58 and 0.68 respectively). Surprisingly, the *direct-only*, where no wave is being generated, had also been mistaken to be a *direct-and-copy or flash-latency* in five out of 28 samples, see Figure 10.

**Figure 10.**
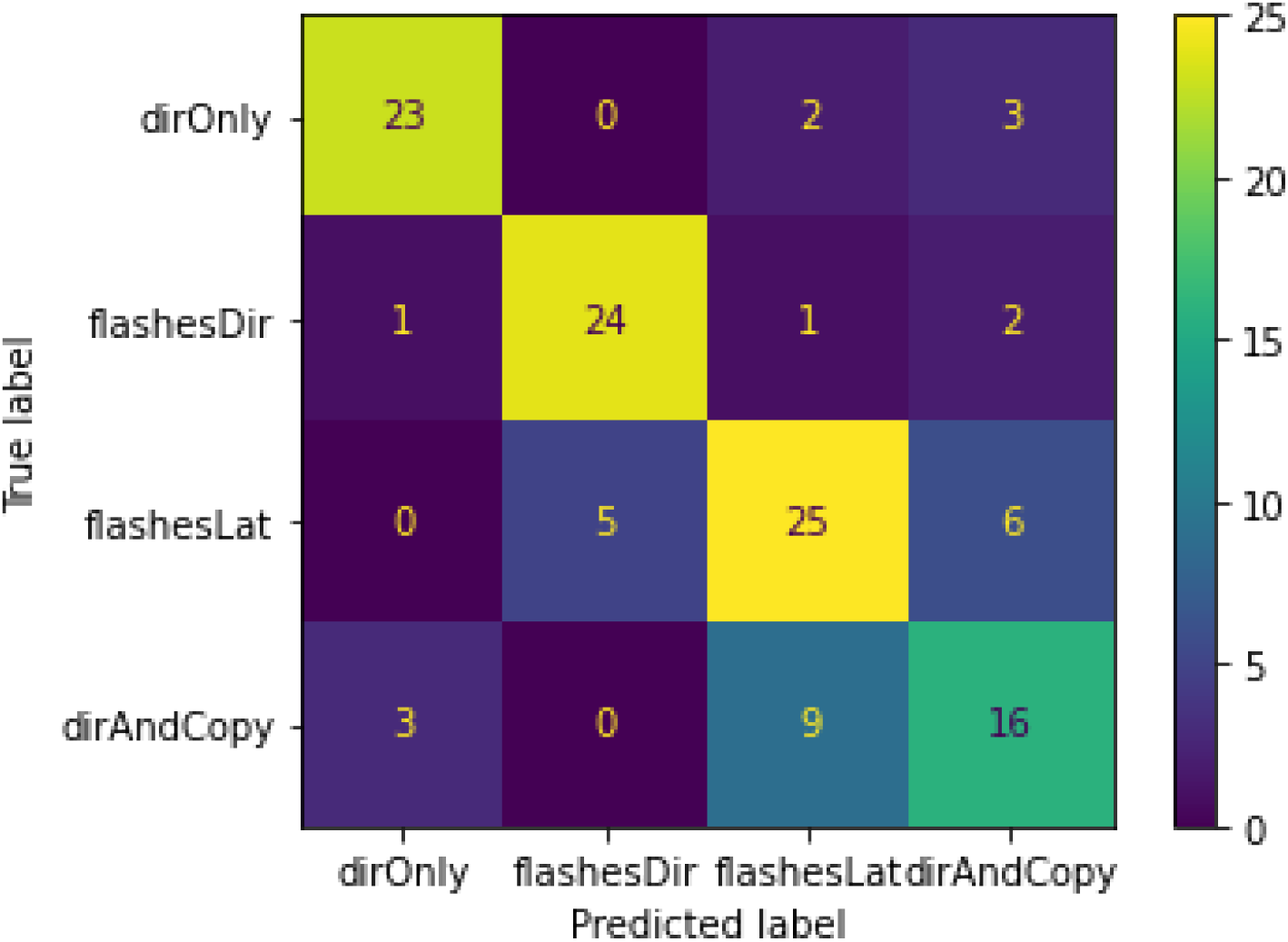
Confusion table of the fish response dataset. The DNN could distinguish among different fish responses. The separation between the direct-and-copy and the flash-latency models is weaker than the others.

## 5. Discussion

The main goal of this work was three-fold. First, to establish whether models of schooling fish can generate realistic flash waves that propagate across the school in response to attack. Second, to demonstrate that these flash patterns contain accessible information relating to the dynamics of the school, the behavior of individuals within it (in particular their response to threats) and to the nature of the attack. Third, to explore the possibility that school members are using this source of information themselves, and test how that would affect the attack-response behavior. Overall, we provide a proof-of-concept which demonstrates that, on the one hand, flash patterns are indeed indicative of the state and dynamics of the school and the behavior of the individuals that compose it; and, on the other hand, that the flashes may be an important causative factor in shaping the escape behavior of the fish.

Models of collective motion typically view the propagation of information as proceeding at a highly localized scale (e.g. (Ballerini et al. 2008; Herbert-Read et al. 2011; Berdahl et al. 2013; Rosenthal et al. 2015)). Our results suggest that light flashes could convey macroscopic, long-range information that may play a role in synchronizing the schooling fish. The high detectability of the flashes facilitates long-distance, many-to-one emergent signaling despite the rapid attenuation of light underwater (i.e., as opposed to the situation in air (Kastberger et al. 2008; Hemelrijk and Hildenbrandt 2011). Because this signal travels instantaneously across long distances (in terms of body lengths) it provides a feasible example for a mechanism that thus far has only been implied (Potts 1984) and alleviates the need to assume that the school is poised near criticality (Bialek et al. 2014).

Our results show that flash-signatures convey enough information to differentiate between schooling scenarios. Event-detection, individual motion rules, and properties of school geometries could all be explored using the flash signature of schools. The abilities demonstrated in this study together with the fact that these flashes are perceivable by the fish, leads to a possibility that a complex and rich flash-language is awaiting to be discovered, which is possibly already in use by the schooling fish and their predators.

## REFERENCES

Achuthan, S., R. Chatterjee, S. Kotnala, A. Mohanty, S. Bhattacharya, R. Salgia, and P. Kulkarni. 2022. Leveraging deep learning algorithms for synthetic data generation to design and analyze biological networks. Journal of Biosciences 47:43.

Aoki, I. 1982. A simulation study on the schooling mechanism in fish. NIPPON SUISAN GAKKAISHI.

Ballerini, M., N. Cabibbo, R. Candelier, A. Cavagna, E. Cisbani, I. Giardina, V. Lecomte, et al. 2008. Interaction ruling animal collective behavior depends on topological rather than metric distance: Evidence from a field study. Proceedings of the National Academy of Sciences 105:1232–1237.

Batty, R. S. 1989. Escape Responses of Herring Larvae to Visual Stimuli. Journal of the Marine Biological Association of the United Kingdom 69:647–654.

Berdahl, A., C. J. Torney, C. C. Ioannou, J. J. Faria, and I. D. Couzin. 2013. Emergent Sensing of Complex Environments by Mobile Animal Groups. Science 339:574–576.

Bialek, W., A. Cavagna, I. Giardina, T. Mora, O. Pohl, E. Silvestri, M. Viale, et al. 2014. Social interactions dominate speed control in poising natural flocks near criticality. Proceedings of the National Academy of Sciences 111:7212–7217.

Brehmer, P., G. Sancho, V. Trygonis, D. Itano, J. Dalen, A. Fuchs, A. Faraj, et al. 2019. Towards an Autonomous Pelagic Observatory: Experiences from Monitoring Fish Communities around Drifting FADs. Thalassas: An International Journal of Marine Sciences 35:177–189.

Brownlee, J. 2019. A Gentle Introduction to the Rectified Linear Unit (ReLU). Machine Learning Mastery.

Couzin, I. D., J. Krause, R. James, G. D. Ruxton, and N. R. Franks. 2002. Collective Memory and Spatial Sorting in Animal Groups. Journal of Theoretical Biology 218:1–11.

Cunningham, P., M. Cord, and S. J. Delany. 2008. Supervised Learning. Pages 21–49 in M. Cord and P. Cunningham, eds. Machine Learning Techniques for Multimedia: Case Studies on Organization and Retrieval, Cognitive Technologies. Springer, Berlin, Heidelberg.

Denton, E. J., and J. a. C. Nicol. 1966. A survey of reflectivity in silvery teleosts. Journal of the Marine Biological Association of the United Kingdom 46:685–722.

Denton, E. J., and D. M. Rowe. 1994. Reflective communication between fish, with special reference to the greater sand eel, Hyperoplus lanceolatus. Philosophical Transactions of the Royal Society of London. Series B: Biological Sciences 344:221–237.

Domenici, P., and R. S. Batty. 1994. Escape manoeuvres of schooling Clupea harengus. Journal of Fish Biology 45:97–110.

Domenici, P., and M. E. Hale. 2019. Escape responses of fish: a review of the diversity in motor control, kinematics and behaviour. Journal of Experimental Biology 222.

Dunn, K. W., C. Fu, D. J. Ho, S. Lee, S. Han, P. Salama, and E. J. Delp. 2019. DeepSynth: Three-dimensional nuclear segmentation of biological images using neural networks trained with synthetic data. Scientific Reports 9:18295.

Gerlotto, F., S. Bertrand, N. Bez, and M. Gutierrez. 2006. Waves of agitation inside anchovy schools observed with multibeam sonar: a way to transmit information in response to predation. ICES Journal of Marine Science 63:1405–1417.

Hemelrijk, C. K., and H. Hildenbrandt. 2011. Some Causes of the Variable Shape of Flocks of Birds. PLOS ONE 6:e22479.

Hemelrijk, C. K., L. van Zuidam, and H. Hildenbrandt. 2015. What underlies waves of agitation in starling flocks. Behavioral Ecology and Sociobiology 69:755–764.

Herbert-Read, J. E., J. Buhl, F. Hu, A. J. W. Ward, and D. J. T. Sumpter. 2015. Initiation and spread of escape waves within animal groups. Royal Society Open Science 2.

Herbert-Read, J. E., A. Perna, R. P. Mann, T. M. Schaerf, D. J. T. Sumpter, and A. J. W. Ward. 2011. Inferring the rules of interaction of shoaling fish. Proceedings of the National Academy of Sciences 108:18726–18731.

Hughey, L. F., A. M. Hein, A. Strandburg-Peshkin, and F. H. Jensen. 2018. Challenges and solutions for studying collective animal behaviour in the wild. Philosophical Transactions of the Royal Society B: Biological Sciences 373:20170005.

Jagtap, H., and M. Chavaan. 2021. Robust Underwater Animal Detection Adopting CNN with LSTM. Pages 195–208 in S. N. Merchant, K. Warhade, and D. Adhikari, eds. Advances in Signal and Data Processing, Lecture Notes in Electrical Engineering. Springer, Singapore.

Jeckel, H., E. Jelli, R. Hartmann, P. K. Singh, R. Mok, J. F. Totz, L. Vidakovic, et al. 2019. Learning the space-time phase diagram of bacterial swarm expansion. Proceedings of the National Academy of Sciences 116:1489–1494.

Kastberger, G., E. Schmelzer, and I. Kranner. 2008. Social Waves in Giant Honeybees Repel Hornets. PLOS ONE 3:e3141.

Lecheval, V., L. Jiang, P. Tichit, C. Sire, C. K. Hemelrijk, and G. Theraulaz. 2018. Social conformity and propagation of information in collective U-turns of fish schools. Proceedings of the Royal Society B: Biological Sciences 285:20180251.

Lee, S.-H. 2006. Predator’s attack-induced phase-like transition in prey flock. Physics Letters A 357:270–274.

Long, J., E. Shelhamer, and T. Darrell. 2015. Fully Convolutional Networks for Semantic Segmentation. Pages 3431–3440 in. Presented at the Proceedings of the IEEE Conference on Computer Vision and Pattern Recognition.

Marras, S., R. S. Batty, and P. Domenici. 2012. Information transfer and antipredator maneuvers in schooling herring. Adaptive Behavior 20:44–56.

Mitash, C., K. E. Bekris, and A. Boularias. 2017. A self-supervised learning system for object detection using physics simulation and multi-view pose estimation. Pages 545–551 in 2017 IEEE/RSJ International Conference on Intelligent Robots and Systems (IROS). Presented at the 2017 IEEE/RSJ International Conference on Intelligent Robots and Systems (IROS).

Pertzelan, A., G. Ariel, and M. Kiflawi. 2022. Light flashes and the geometry of specular fish schools. Journal of The Royal Society Interface 19:20210906.

Potts, W. K. 1984. The chorus-line hypothesis of manoeuvre coordination in avian flocks. Nature 309:344–345.

Radakov, D. 1973. Schooling in the ecology of fish.

Raghunathan, T. E. 2021. Synthetic Data. Annual Review of Statistics and Its Application 8:129–140.

Reynolds. 1987. Flocks, herds and schools: A distributed behavioral model | Proceedings of the 14th annual conference on Computer graphics and interactive techniques.

Rosenthal, S. B., C. R. Twomey, A. T. Hartnett, H. S. Wu, and I. D. Couzin. 2015. Revealing the hidden networks of interaction in mobile animal groups allows prediction of complex behavioral contagion. Proceedings of the National Academy of Sciences 112:4690–4695.

Shi, X., Z. Chen, H. Wang, D.-Y. Yeung, W. Wong, and W. Woo. 2015. Convolutional LSTM Network: A Machine Learning Approach for Precipitation Nowcasting. Advances in Neural Information Processing Systems (Vol. 28). Curran Associates, Inc.

Sonoda, K., H. Murakami, T. Niizato, T. Tomaru, Y. Nishiyama, and Y.-P. Gunji. 2019. Propagating wave based on transition of interaction within animal group. Biosystems 185:104019.

Treherne, J. E., and W. A. Foster. 1981. Group transmission of predator avoidance behaviour in a marine insect: The trafalgar effect. Animal Behaviour 29:911–917.

Vicsek, T., and A. Zafeiris. 2012. Collective motion. Physics Reports, Collective motion 517:71–140.

Xu, Y., and R. Goodacre. 2018. On Splitting Training and Validation Set: A Comparative Study of Cross-Validation, Bootstrap and Systematic Sampling for Estimating the Generalization Performance of Supervised Learning. Journal of Analysis and Testing 2:249–262.

Yin, X., D. Wu, Y. Shang, B. Jiang, and H. Song. 2020. Using an EfficientNet-LSTM for the recognition of single Cow’s motion behaviours in a complicated environment. Computers and Electronics in Agriculture 177:105707.

